# Genotype-by-environment-by-environment (G×E×E) interactions have the potential to shape adaptation in response to multiple stressors

**DOI:** 10.64898/2026.07.21.739626

**Authors:** Sarah E. Britton, Alfonso Ruiz Quintero, Sanam Rozycki-Shah, Patrick T. Rohner

**Author notes:** Corresponding author: SEB.

## Abstract

Rapid adaptation in complex environments depends not only on the amount of genetic variation, but also on patterns of covariation among traits targeted by selection. Anthropogenic stressors create rapidly changing and multifaceted environments and provide powerful systems in which to investigate the potential for adaptation to multiple coinciding stressors. We investigate the combined effects of heat and chemical stress on survival in the black scavenger fly, *Sepsis neocynipsea*, and determine the genetic basis for resistance. In a fully factorial experiment, we expose isofemale lines to combinations of heat stress and ivermectin, a veterinary antiparasitic to which these flies are naturally exposed in agricultural landscapes. Using a Bayesian quantitative genetic approach, we estimate broad-sense genetic variation and cross-environmental genetic correlations. First, we show that these two stressors have synergistic effects on survival, with heat stress exacerbating the lethal effects of ivermectin. Second, we find that the largest component of genetic variation is the response to heat and ivermectin in combination (genotype-by-environment-by-environment; G×E×E). Third, cross-environmental genetic correlations are weak, implying that relative genetic performance is dependent on the specific combination of stressors. Together, these results suggest that incorporating G×E×E is essential for understanding adaptive potential in multi-stressor environments.

## Introduction

Adapting rapidly to new selective pressures depends heavily on the amount of standing genetic variation (Barrett & Schluter, 2008; Bitter et al., 2019; Chaturvedi et al., 2021).

However, because natural selection acts on multiple co-varying traits simultaneously, evolution depends not only on variance in individual traits but also on the genetic architecture of their covariance (A. F. Agrawal & Stinchcombe, 2009; Via, 1991; Via & Lande, 1985). This is particularly true in complex environments where selective pressures from multiple environmental factors are acting concurrently. Anthropogenic stressors, such as climate change, urbanization, pollutants, and introduced species, are a unique challenge for organisms because they are often novel and rapidly changing. Critically, these new stressors are not faced in isolation but are often experienced in combination. Thus, it is important to understand not only how these stressors may interact non-additively to influence the behavior, physiology, life history, and ultimately fitness of susceptible organisms, but also the potential for adaptation in these novel and complex environments.

The combination of environmental stressors on organismal performance and survival may combine additively, such that effects are a result of both stressors acting independently (e.g., Gieswein et al., 2017). However, when stressors combine synergistically, their collective impact on organisms is more than the additive effects alone (e.g., Kelly et al., 2016). Alternatively, when stressors combine antagonistically their total effects on an organism are less than the additive effects, such that the presence of one stressor mitigates the effects of another (e.g., Rodgers & Gomez Isaza, 2023). This ‘cross-resistance’ or ‘cross-protection’ is commonly observed and may be explained by shared response pathways or protective mechanisms that can alleviate stress from multiple sources, such as upregulation of heat shock proteins or reactive oxygen species (ROS) signaling (Bueno et al., 2023; Rodgers et al., 2025).

While the phenotypic consequences of interacting stressors is now well-researched (Bueno et al., 2023; Coors & De Meester, 2008; Gunderson et al., 2022; Kaunisto et al., 2016; Todgham & Stillman, 2013), it is much less clear whether genetic variation exists for responses to these interactions, limiting our ability to predict evolution in complex environments. Genetic variation in stress resistance is common (e.g., Pereira et al., 2017), and resistance to one stressor may genetically covary with resistance to another, either negatively (e.g., Olsen et al., 2019), potentially slowing adaptation, or positively (e.g., Bubliy & Loeschcke, 2005), potentially facilitating adaptation. Stress resistance can be measured as performance (e.g., survival, growth) across an environmental range (i.e., less stressful to more stressful) and genetic variation can be quantified by measuring genotype-by-environment (G×E) interactions, ,(i.e., how genotypes differ in their response to environments). This reaction norm approach (Schlichting & Pigliucci, 1998) makes it feasible to disentangle genotypes that are superior regardless of the stressor, versus those that are particularly tolerant of a certain stressful environment. However, the genetic response to one stressor may also depend on other environmental variables, which can be captured by quantifying genotype-by-environment-by-environment (G×E×E) interactions, or genetic variance in response to a combination of environmental factors. These higher-order interactions are often overlooked but potentially crucial to understand and predict evolutionary responses in complex environments (Schmidlin et al., 2024). If there is genetic variation in response to combined stressors (G×E×E; i.e., genetic variation for cross-resistance or trade-offs), then selection will favor those genotypes with the fittest response to both pressures, which may differ from those genotypes that perform best in response to a single pressure. This may explain why stress tolerance traits do not always evolve as anticipated in response to selection, either in the lab (Kelly et al., 2016) or across environmental clines (González-Tokman et al., 2022).

In this study we consider the effects of two environmental stressors: heat and chemical exposure. The widespread effects of climate change often result in more extreme heat, both in terms of increasing mean temperatures and short-term extreme heat events (Rahmstorf & Coumou, 2011; Wigley, 2009). There is a large literature on how organisms cope with heat stress (Couper et al., 2025; Kelly et al., 2012; Ma et al., 2020), and while some populations harbor variation in heat tolerance, others have limited variation which may slow adaptation (Fallis et al., 2011; Nati et al., 2021). Chemical stressors in the form of pesticides or industrial and urban run-off are another major source of environmental change and often introduce challenges that are novel to organisms in their environment. On agricultural landscapes pharmaceuticals administered to animals can leach into the environment. One pharmaceutical of concern is ivermectin, a broad-spectrum antiparasitic used to treat livestock infected with parasites including nematodes, lice, ticks, mites and flies (Crump, 2017). The use of ivermectin is widespread in the United States and up to 90% of the administered drug can be excreted unmetabolized into the environment, lasting for weeks or months (Alvinerie et al., 1999; Herd et al., 1996). By targeting chloride channels, this drug induces neuro-muscular paralysis and has widespread lethal effects on target as well as on many non-target arthropods (Jochmann & Blanckenhorn, 2016; Puniamoorthy et al., 2014; Sánchez-Bayo, 2012). Of particular concern are dung-feeding insects which are not only highly sensitive to ivermectin, but also directly exposed to unmetabolized ivermectin in livestock dung. Because these insects provide important ecosystem services in agricultural environments (Anderson et al., 2024; Hajji et al., 2024; Nichols et al., 2008; Tixier et al., 2015), understanding their response to these chemicals is important for management.

Here, we use the black scavenger fly, *Sepsis neocynipsea* (Diptera: Sepsidae), to investigate the combined effects of heat and ivermectin stress on survival and genetic variation in survival, and the potential for adaptation in response to these stressors. *S. neocynipsea* is widespread across the United States and Europe and feeds on vertebrate dung as both larvae and adults (Pont & Meier, 2002). Females lay their eggs into dung pads where larvae develop and pupate. Although adults are primarily nectar-feeders, dung is an important source of protein, especially for ovipositing females. Previous work shows that this and related species are vulnerable to ivermectin in both larval and adult stages (Blanckenhorn et al., 2013; Conforti et al., 2018; Puniamoorthy et al., 2014; Walker et al., 2025). In addition, sepsid flies are also sensitive to heat stress; in this and related species heat stress depends on life stage, population origin, acclimation experience, and the trait in question. For instance, survival during the larval stage drops quickly at temperatures above 32°C (Khelifa et al., 2019), however, adults of some species are still active at temperatures up to 45°C, although it is unclear how long this is sustainable (Kjærsgaard et al., 2023). Evidence suggests that heat and chemical stressors may combine non-additively in their effect on insects. Heat stress can lower the ability of insects to combat pesticides due to increasing metabolic costs, higher uptake rates, increased membrane permeability, and smaller body sizes (Dinh Van et al., 2014; Noyes et al., 2009; Verheyen et al., 2022). On the other hand, heat can offer cross-protection rendering insecticides less lethal, especially via the upregulation of heat shock proteins (Gu et al., 2010; Robertson, 2004; Sørensen et al., 2003). Here, we test how these stressors interact and how they impact genetic variation.

We first ask whether the combined effects of heat and ivermectin exposure on survival are additive or interactive in this species. Then, we take a quantitative genetics approach to measure genetic variation in resistance to heat and ivermectin alone, as well as in combination, to test whether G×E×E is a significant source of variation. Finally, we investigate the genetic architecture of these responses, whether resistance to these stressors co-varies, and how genetic responses correlate across environments. These insights will allow us to understand how organisms in complex, multi-stressor environments respond to these anthropogenic changes in the short term and the potential for longer term adaptation.

## Methods

### Isofemale lines

Wild *S. neocynipsea* females were caught by net or aspiration on or near cow dung in June 2025 on open rangeland in Santa Ysabel, California. Individuals were brought back to the lab to confirm sex and species and housed in individual containers provided with sugar, water, and cow dung on which to lay eggs. Offspring of successful females were kept separately as isofemale lines and maintained at 21°C with ad libitum sugar, water, and cow dung until the experiment (4-5 generations, with more than 50 individuals maintained in the culture once established).

### Experiment

Individuals from 49 isofemale lines across four experimental blocks were used for the experiment. For each block, individuals from within a line were divided into four treatments in a 2×2 factorial design. These individuals were collected by adding fresh dung trays to the cultures, allowing females to lay eggs for 48 hours, isolating the trays for development, and capturing emerging the adults. The treatments included two temperature treatments, control (19°C) and heat stress (36°C) and two ivermectin treatments, control and ivermectin stress. Heat stress temperature and ivermectin concentration were chosen to be ecologically relevant and were based on preliminary trials to ensure that they were stressful (i.e, mortality rates differed from control), but not so stressful as to result in complete mortality (and thus, no phenotypic or genetic variation to measure). Flies in the ivermectin stress treatment were given dung with an ivermectin concentration of 750 µg/kg, which was made by first dissolving ivermectin into acetone and then mixing the acetone solution into the dung. Dung was left uncovered overnight to allow for acetone evaporation and then mixed well. To control for any effects caused by using acetone as a solvent, the same amount of acetone was added to the control dung. During the experiment flies were housed in narrow *Drosophila* vials (9.5 × 2.5 cm) with cellulose acetate stoppers (Genesee Scientific, CA). Within each treatment approximately 10 (range = 7-12) mature flies (5-7 days old) were transferred into a vial with sugar and 3g cow dung (either control or ivermectin). Dung and sugar were kept in plastic dishes placed in the vials (4×2 cm and 2×2 cm, respectively). Vials were then transferred into an incubator (Insect Growth Chamber, Caron Scientific, OH) set at 19°C or 36°C for five days. Photoperiod in the incubator was set at 12 light:12 dark. After five days all flies were collected and scored for survival.

Each of the 49 lines had 2-3 replicates per each of the four treatments resulting in 4,637 total flies. Based on previous work in this species (Walker et al., 2025) we expected that sex and size would be important covariates and might have complex interactions with the main treatment variables. We therefore determined the sex of each individual based on the presence of male-specific spikes on the forefemur and measured the length of the hindtibia as an overall estimate of body size (Rohner, 2021). Sex and size measurements were split randomly and conducted by two experimenters who were consistent in their measurements of length (r^2^ = 0.89, CV=1.8%, n=15).

### Statistical methods

We took a multistep approach to model the probability of survival with temperature treatment, ivermectin treatment, sex, and size (and their interactions) as fixed effects and line and block as random effects. In the following models hindtibia length was centered, but no other transformations were necessary.

#### Determining fixed effect structure

Because we expected complex interactions between fixed factors, we first determined the fixed effect structure using maximum likelihood techniques. Using the lme4 package (Bates et al., 2015), we built generalized mixed effect models with a binomial distribution and line and block as random factors, and then followed standard backwards model elimination selection protocol. We began with the most complex model, which included a 4-way interaction, then proceeded with stepwise model simplification until a final model containing only one significant 3-way interaction remained. We compared models via both AIC and log ratio tests (Tables S1 and S2). The most suitable model was as follows:

Probability of Survival = Temperature + Ivermectin + Size + Sex + (Temperature × Ivermectin) + (Temperature × Size) + (Ivermectin × Size) + (Ivermectin × Sex) + (Sex × Size) + (Temperature × Ivermectin × Size)

#### Bayesian modeling

Due to complete data separation, which is common in complex designs with binary response variables, it was not possible to estimate variance components using standard maximum likelihood approaches. We therefore used a Bayesian approach to estimate genetic effects. Specifically, based on the fixed effects structure determined above, we used the brms package (Bürkner, 2017) to test the explanatory power of different random effect structures. We fit all models with a Bernoulli distribution and a logit link. Models were run with four chains for 4,000 iterations and 2,000 burn-ins each. We set weakly informative priors for both fixed and random effects (Table S3). We confirmed model convergence by checking that the chains were sufficiently mixed. Models were compared with a leave-one-out procedure using the loo package (Vehtari et al., 2021) to extract the estimate of the expected log predictive density (ELPD). The loo package was also used to compare ELPD values to determine the best supported model, with a lower ELPD signifying a better fit. For the hierarchical and Bernoulli models that we used, ELPD is a superior model comparison method to others such as WAIC (Vehtari et al., 2021).

We tested and compared the following random effects structures in our models:

1. Null: Random effect of block only
2. G: Random effects of block and line
3. G×E (temperature only): Random effects of block, line, and line × temperature
4. G×E (ivermectin only): Random effects of block, line, and line × ivermectin
5. G×E (both): Random effects of block, line, line × temperature, and line × ivermectin
6. G×E×E: Random effects of block, line, line × temperature, line × ivermectin, and line × temperature × ivermectin

#### Genetic variance

We extracted broad sense genetic variance distributions from the best supported model by calculating variance as twice the among line variance component (2×s.d.^2^) because the isofemales lines are treated as full-sibling families (David et al., 2005; Hoffmann & Parsons, 1988). We also extracted breeding value distributions from the best supported model and estimated posterior mean breeding values. These line-specific values represent relative genetic deviations in performance for a specific environment. If there are no G×E interactions we expect breeding values for specific lines to be consistent across environments (i.e., the same lines are always performing better or worse compared to others.) However, with G×E interactions we expect to see a re-ordering of breeding values across environments.

#### Cross-environmental genetic correlations

To understand the structure of genetic correlations between stress responses, we quantified cross-environmental genetic correlations by re-running the brms model using the full environment (temperature and ivermectin treatment combination) as a factor with four levels, with survival in each environment modeled as a separate trait. We modeled environment-specific values by fitting line as a random effect that interacts with environment, allowing for an unstructured correlation among environments, and extracted the distribution of correlations. If there is no G×E×E, we expect high correlations (i.e., genetic performance is similar across all environments), while low or negative correlations would indicate G×E×E (i.e., genetic performance depends on the specific environmental combination). For example, this pattern would signify that relative heat tolerance depends on presence of ivermectin (and vice versa).

## Results

### Main treatment effects

The following results are presented as mean estimates with 95% credible intervals. Heat stress and ivermectin stress both have negative effects on survival (heat stress: estimate = -2.20 [-2.76, -1.67]; ivermectin stress: estimate = -4.11 [-4.72, -3.54]; Fig. 1, Table S4). Furthermore, these stressors have a synergistic effect whereby survival is worse than expected when exposed to both (temperature x ivermectin: estimate = -1.42 [-2.09, -0.75]; Fig. 1, Table S4). Sex significantly interacted with ivermectin such that ivermectin stress impacted females more than males (ivermectin x sex: estimate = 2.17 [1.77, 2.57; Fig. 1a, c, Table S4).

**Figure 1.**
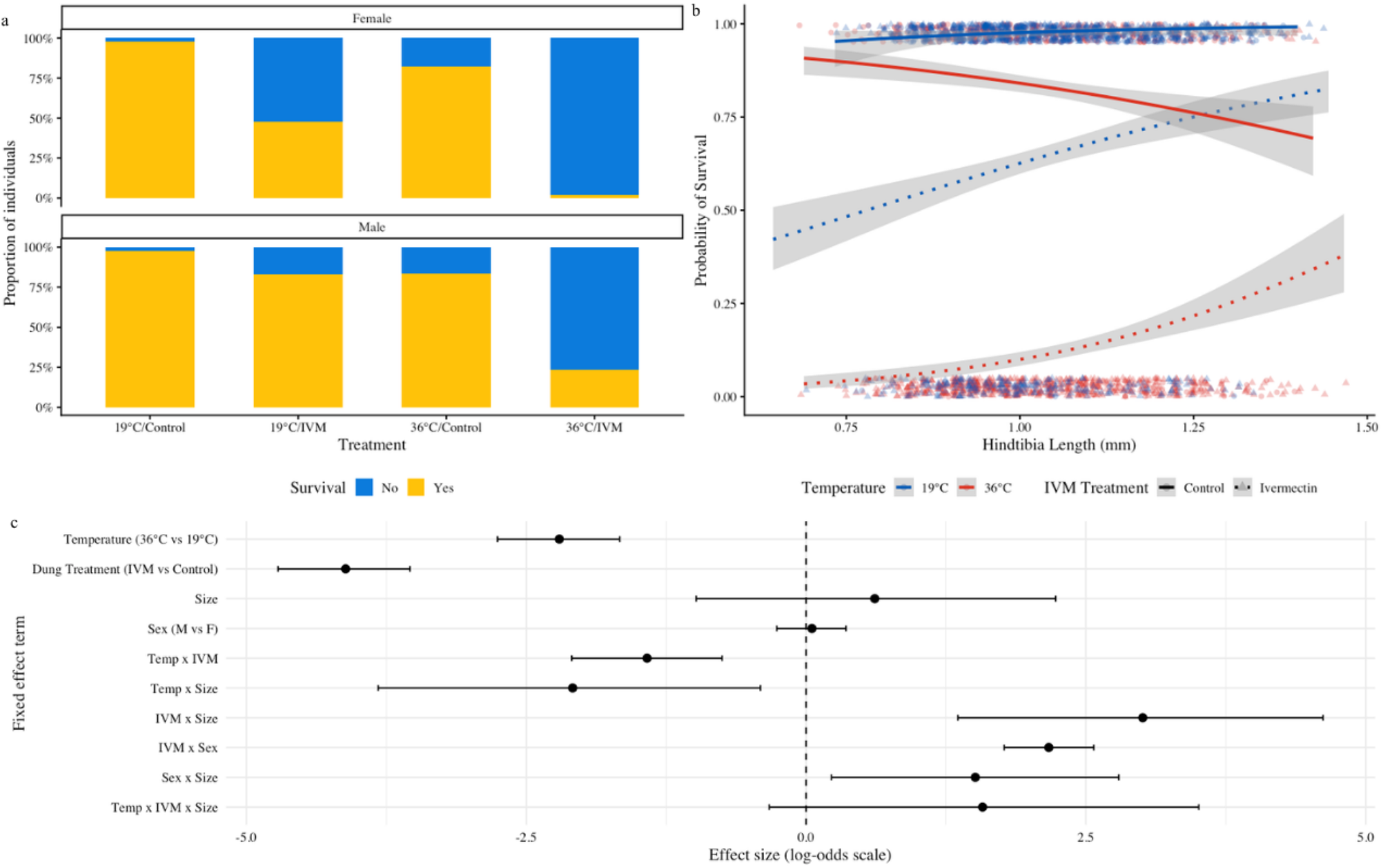
Effects of fixed factors on survival. (a) Raw data showing proportion of surviving individuals as a function of treatment (temperature and ivermectin) and sex. (b) Raw data showing probability of survival as a function of size (hindtibia length), temperature, and ivermectin treatment. (c) Effect sizes and 95% credible intervals of the fixed effects from the best supported model.

Size significantly interacted with ivermectin such that small individuals were less likely to survive when exposed to ivermectin (ivermectin x size: estimate = 3.01 [1.36, 4.62]; Fig. 1b, c, Table S4). Size also significantly interacted with temperature such that large individuals were less likely to survive under heat stress (temperature x size: estimate = -2.09 [-3.82, -0.41]; Fig 1b, c).

### Genetic variance

The full model including G×E×E had the best overall support (Table 1). Importantly, the G×E×E component was also the largest component of genetic variance, indicating that lines differ in how they respond to the interaction of heat and ivermectin stressors (Fig. 2a, Table S5). Additionally, breeding values show re-ordering across environments, consistent with the presence of both G×E and G×E×E interactions (Fig. 2b). Note that due to our experimental design genetic variance is broad-sense and we cannot separate effects from dominance, maternal effects, or epistatic components.

**Table 1.** Model comparison based on leave-one-out procedures for different random effect structures. Models are compared with expected log predictive density (ELPD) values.

| Model | ELPD | Change ELPD |
| --- | --- | --- |
| Null | -1687.73 | -109.5 |
| G | -1622.35 | -44.1 |
| G×E (temperature) | -1607.79 | -29.5 |
| G×E (IVM) | -1595.61 | -17.3 |
| G×E (both) | -1590.31 | -12.0 |
| G×E×E | -1578.27 | 0 |

**Figure 2.**
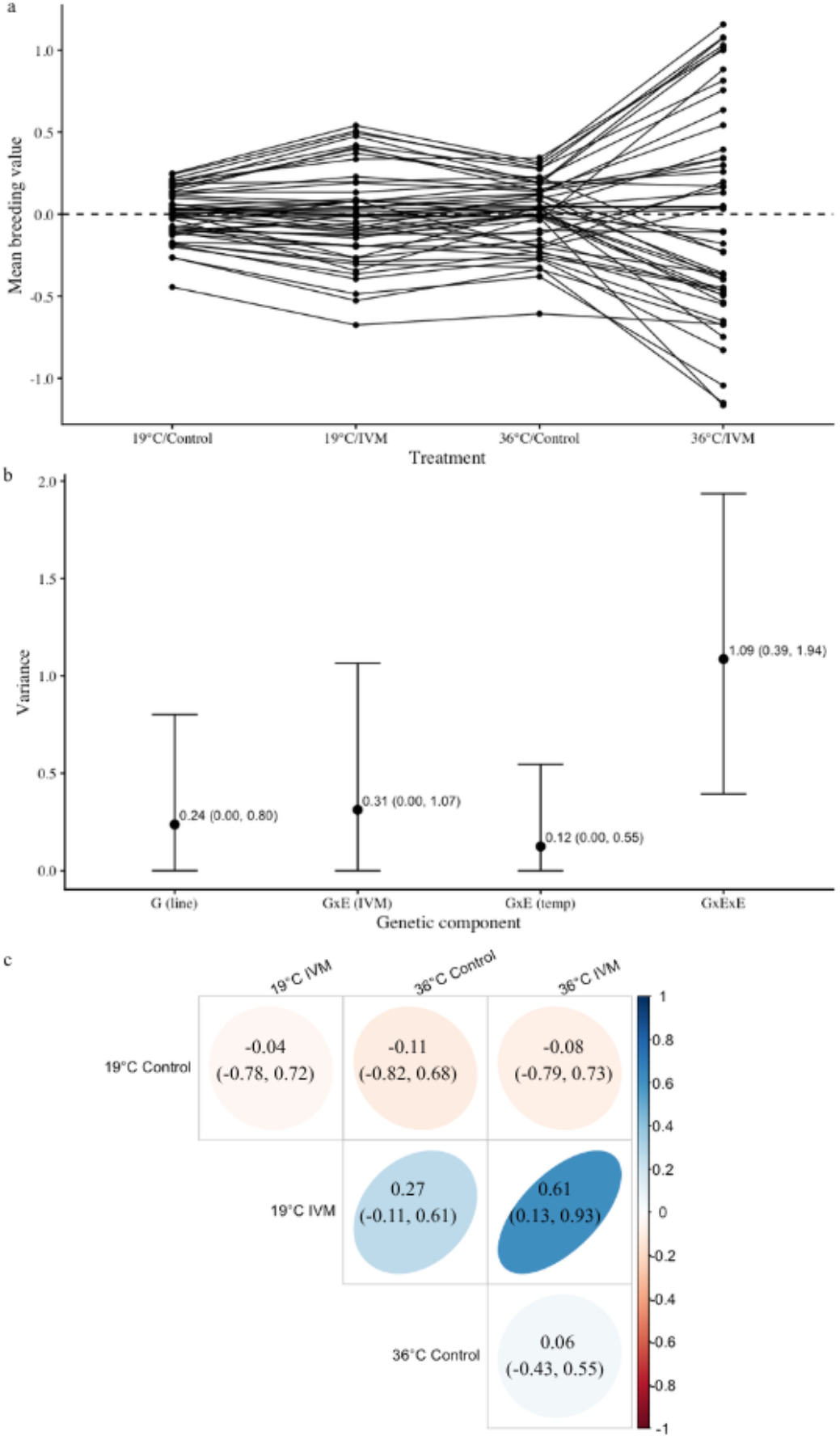
Broad-sense genetic variance and cross-environmental correlations. (a) Distribution of variance across components calculated from the best-supported model with mean and 95% credible intervals (block and residual variance not shown). (b) Posterior mean breeding values (relative genetic performance) for each line shown as reaction norms across combinations of temperature and ivermectin treatments. Positive values are above the population mean while negative values below are below the population mean. Points represent mean estimates for a line and are shown without 95% credible intervals for clarity. (c) Cross-environmental genetic correlations. Darker blue colors represent more positive correlations while darker red colors represent more negative correlations. Rounder ellipses represent weaker correlations. Mean correlation value with 95% credible intervals are shown for each comparison.

### Cross-environmental correlations

In general, cross-environmental genetic correlations were non-significant (95% credible intervals overlap 0), signifying that relative performance of a particular line in one environment shows no strong correspondence to its performance in another environment (Fig. 2c, Fig. S1). However, for some of the correlations 95% credible intervals are quite wide, indicating high uncertainty in the parameter estimates. Consistent with these findings we found that breeding values are not constant across environments, but rather that relative genetic performance is dependent on the specific treatment combination (Fig. 2b).

To investigate the genetic correlation between stress responses, we measured the correlation in performance between the 36°C/Control environment (heat tolerance) and the 19 °C/IVM environment (ivermectin resistance). We found a non-significant correlation (r^2^ = 0.27 [-0.11, 0.61]), indicating little genetic relationship in how flies respond to these two stressors. This is not consistent with evidence for either a genetic trade-off between stress responses, or variation for cross-resistance. To further investigate the genetic relationship between ivermectin resistance and heat tolerance, we measured the correlation in performance between the 36°C/Control environment and the 36°C/IVM environment. Again, we found a non-significant correlation (r^2^ = 0.06 [-0.43, 0.55]), which implies that relative line performance in response to temperature stress depends on the ivermectin treatment. We also considered the reverse relationship by measuring the correlation in performance between the 19°C/IVM environment and the 36°C/IVM environment. Here, we found a significant positive correlation (r^2^ = 0.61 [0.13, 0.93]), indicating that some lines were performing better than others when exposed to ivermectin, regardless of temperature.

## Discussion

In complex environments where organisms face many selective pressures simultaneously, G×E×E interactions may dictate the direction and speed of adaptation because traits can co-vary in ways that can either hinder or facilitate adaptation (A. F. Agrawal & Stinchcombe, 2009). This is especially true for anthropogenic stressors, many of which are novel, rapidly changing, and can combine unpredictably. It is thus critical to our understanding of adaptation in rapidly changing environments how organisms respond to stressors alone and in combination, and how genetic variance for response to these stressors is structured. Here we address this problem by investigating how a population of *S. neocynipsea* flies responds to the combined effects of heat stress and ivermectin, using a quantitative genetics approach. First, we find that heat stress greatly exacerbates the effects of ivermectin on survival. Second, we show that genetic variation for stressor interactions is a major component of broad-sense genetic variance. Finally, we find limited evidence for strong genetic correlations among stress responses, indicating that genetic values may be difficult to predict across environments. These findings highlight the potential for G×E×E interactions to shape an adaptive response.

### Synergistic effects of heat and ivermectin on survival

Exposure to both heat stress and ivermectin led to lower survival for *S. neocynipsea* flies, with exposure to both stressors exacerbating this effect, in line with the predictions for a synergistic response. This is consistent with other insect studies investigating the combined effects of heat and chemical stressors (González-Tokman et al., 2022; Huisamen et al., 2023). Higher metabolic rates in warm environments can increase oxidative damage and the need for protective antioxidants (Chen et al., 2018; Jena et al., 2013), which may then compete for antioxidants required for an ivermectin response. However, heat stress has also been found to increase ivermectin resistance in some insects (Esquivel-Román et al., 2025; Walker et al., 2025), potentially due to the upregulation of heat shock proteins, which are also used in an ivermectin response (Bueno et al., 2023; Villada-Bedoya et al., 2021). Whether these stressors combine synergistically or antagonistically may depend on the degree of heat stress (i.e., the length of exposure and proximity to the thermal maximum). Milder stressors may be more likely to have hormetic effects, such that a milder heat stress may be more likely to help organisms cope with chemical stressors like ivermectin. For instance, previous work in this species found that milder heat stress (33°C), which did not alone impact survival, combined antagonistically with ivermectin, leading to a non-additive increase in survival when combined (Walker et al., 2025).

Size is an important moderator of these stressors as larger flies fared better than smaller flies, specifically in the presence of ivermectin (Fig. 1), consistent with previous work in this system (Walker et al., 2025). This is not surprising as larger insects are often more robust to chemical stress (Gergs et al., 2015). Sex was also an important modulator as males fared better than females in ivermectin treatments (Fig. 1), consistent with previous work (Walker et al., 2025). The reason for this is unclear, especially because many insect studies document that females are more robust to chemical toxins than males (Foley et al., 2019; Lazarević et al., 2022; Liu et al., 2025). One explanation is behavioral differences wherein females may have faced stronger ivermectin exposure if they were more likely to feed on dung than males due to higher protein needs for egg development.

### G×E×E interactions are a major contributor to broad-sense genetic variance

To understand adaptive potential in multi-stressor environments we tested for genetic variation in response to heat stress, ivermectin stress, and their combination (G×E×E). We find that G×E×E is the largest component of genetic variance (Fig. 2a). In other words, genetic lines in this population vary not only in how they respond to these stressors individually but also how they respond to them in combination. This is clearly illustrated by examining the breeding values of genetic lines across environments (Fig. 2b), which shows that relative line performance depends on the specific combination of stressors. Similar patterns were found in the yellow dung fly, where larval growth and development showed heritable variation in response to the combination of heat and ivermectin stress (González-Tokman et al., 2022). It is important to note that our measures of genetic variation are broad-sense and thus we cannot quantify the degree of variation explained by dominance, epistasis, or maternal effects. Despite these limitations, the data are consistent with the hypothesis that G×E×E might be an underappreciated component of genetic variation that has been overlooked as a determinant of adaptation in complex environments.

### Limited evidence for trade-offs and cross-environmental correlations

To get a better understanding of the nature of G×E×E we quantified cross-environmental correlations (Fig. 2c). High positive correlation values indicate that relative performance (line survival) is similar across environments while a high negative correlation value indicates a genetic performance trade-off across environments. First, we were interested in comparing line performance in the heat stress only (36°C/ Control) and ivermectin stress only (19°C/ IVM) environments because this correlation can reveal genetic trade-offs between the response to these two stressors individually. Genetic trade-offs are thought to be important constraints in limiting, or at least slowing, adaptation (A. A. Agrawal et al., 2010; A. F. Agrawal & Stinchcombe, 2009; Connallon & Hall, 2018). We find no strong evidence for a correlation between how genetic lines respond to heat stress and ivermectin stress. Thus, while we do not document a trade-off that may constrain evolution, we also do not find strong evidence for facilitation.

Next, we were interested in how relative performance of genetic lines in response to one stressor depends on the level of another. In other words, does relative performance in an ivermectin environment depend on temperature (and vice versa)? Without G×E×E we expect high correlations, while low or negative correlations indicate G×E×E, because how well a line performs in a stressful environment depends on the level of the other stressor. We find that the relative performance of genetic lines when exposed to ivermectin remains relatively unchanged across both temperature treatments (high genetic correlation correlation), indicating that some genetic lines are superior at tolerating ivermectin, and suggests that this population may be able to evolve resistance to ivermectin regardless of selective pressure from temperature. However, the same pattern did not hold for the effect of heat stress across ivermectin treatments; we found that relative performance of genetic lines in response to heat depends on whether they are exposed to ivermectin. Predicting evolutionary responses in complex environments may thus require taking G×E×E into account. Incorporating these interactions may greatly enhance our ability to understand and predict evolutionary responses in anthropogenic environments.

However, the degree of genetic variation for cross-resistance or trade-offs across a wider range of traits and taxa remains to be investigated.

It should be noted that many of our measures of genetic correlation have quite wide credible intervals, indicating uncertainty, although the specific correlations in which we were most interested had higher densities (confidence) around the mean (Fig. S1). It is possible that an experimental design with more isofemale line replicates would help clarify these patterns, although we were still able to detect one significant non-zero correlation.

## Conclusions

Standing genetic variation is crucial for adaptive responses, especially on short time scales, such as those imposed by recent anthropogenic stressors. While the importance of this is well-understood, what is often overlooked is how genetic variation and the prognosis for adaptation may change when considering multiple stressors simultaneously. Our findings are consistent with a significant contribution of G×E×E interactions in structuring genetic values as well as genetic variance. Understanding the contribution of G×E×E to the speed and direction of adaptation is important because it implies that selection for genotypes that perform best under a single stressor is not equivalent to selecting for the best genotypes under combined stressors.

Thus, considering the complexity of the selective environment may change the predictions for the strength and direction of selection in the face of multiple stressors, especially in novel anthropogenic environments.

