## Supplemental materials for "Genotype-by-environment-by-environment (G×E×E) interactions have the potential to shape adaptation in response to multiple stressors"

Table S1. AIC and log ratio test (LRT) results for maximum likelihood model comparisons. The reduced 3-way interaction model with only one significant 3-way interaction was used. *Justification for more complex model.

| Model | AIC | LRT |
|---|---|---|
| 4-way interaction | 3295.5 | |
| 3-way interactions (all) | 3294.6 | p = 0.30 |
| *3-way interactions (reduced) | 3295.6 | p = 0.06 |
| 2-way interactions (all) | 3296.8 | p <0.001* |

Table S2. Fixed effects results for final maximum likelihood model.

| Term (baseline) | Estimate | p-value |
|---|---|---|
| Temperature (36) | -2.28 | <0.001 |
| Ivermectin (IVM) | -4.15 | <0.001 |
| Sex (M) | 0.02 | 0.866 |
| Size | 2.30 | 0.183 |
| Temperature x Ivermectin | -1.20 | <0.001 |
| Ivermectin x Sex | 2.13 | <0.001 |
| Ivermectin x Size | 1.06 | 0.549 |
| Temperature x Size | -4.34 | 0.015 |
| Sex x Size | 1.72 | 0.016 |
| Temperature x Ivermectin x Size | 4.06 | 0.043 |

Table S3. Weakly informative priors used for brms models.

| Class | Distribution | Parameters |
|---|---|---|
| b (fixed effects) | Normal | (0, 1.5) |
| Intercept | Student's t | (4, 0, 4) |
| sd (random effects) | Exponential | 1.5 |

Table S4. Estimates of fixed effects from the final Bayesian model with 95% credible intervals.

| Term (baseline) | Estimate | 95% CI |
|---|---|---|
| Temperature (36) | -2.20 | (-2.76, -1.67) |
| Ivermectin (IVM) | -4.11 | (-4.72, -3.54) |
| Sex (M) | 0.05 | (-0.26, 0.36) |
| Size | 0.61 | (-0.98, 2.23) |
| Temperature x Ivermectin | -1.42 | (-2.09, -0.75) |
| Ivermectin x Sex | 2.17 | (1.77, 2.57) |
| Ivermectin x Size | 3.01 | (1.36, 4.62) |
| Temperature x Size | -2.09 | (-3.82, -0.41) |
| Sex x Size | 1.51 | (0.23, 2.79) |
| Temperature x Ivermectin x Size | 1.58 | (-0.33, 3.51) |

Table S5. Estimates of random effects from the final Bayesian model.

| Term | Estimate | 95% CI |
|---|---|---|
| Block | 0.59 | (0.19, 1.43) |
| Line | 0.30 | (0.02, 0.63) |
| Line x Temperature | 0.20 | (0.01, 0.52) |
| Line x Ivermectin | 0.34 | (0.01, 0.73) |
| Line x Temperature x Ivermectin | 0.72 | (0.44, 0.98) |

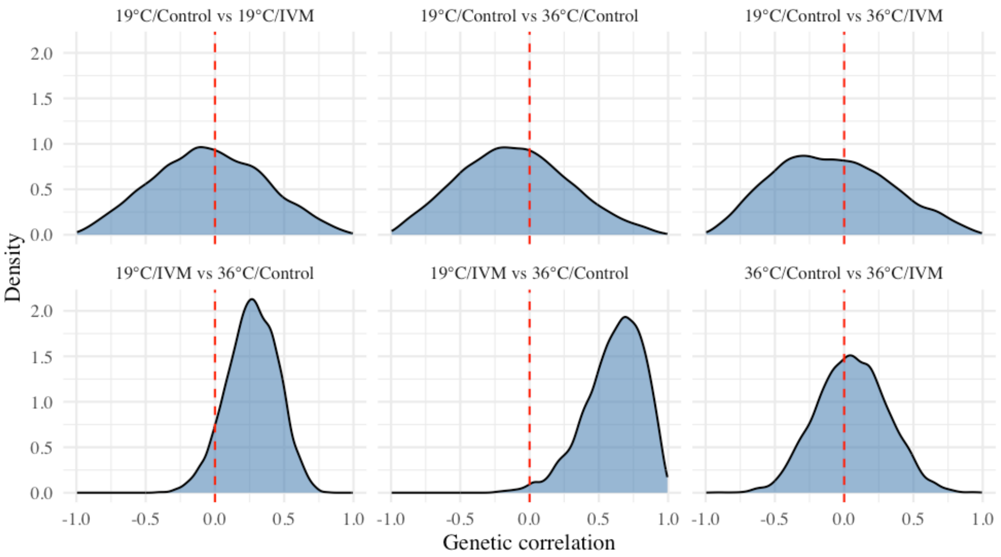


Figure S1. Density plots of correlation values for pairwise cross-environmental genetic correlations.
